# CD32^+^ and PD-1^+^ Lymph Node CD4 T Cells Support Persistent HIV-1 Transcription in Treated Aviremic Individuals

**DOI:** 10.1101/329938

**Authors:** Alessandra Noto, Francesco A. Procopio, Riddhima Banga, Madeleine Suffiotti, Jean-Marc Corpataux, Matthias Cavassini, Craig Fenwick, Raphael Gottardo, Matthieu Perreau, Giuseppe Pantaleo

## Abstract

A recent study conducted in blood has proposed CD32 as the marker identifying the ‘elusive’ HIV reservoir. We have investigated the distribution of CD32^+^ CD4 T cells in blood and lymph nodes(LNs) of healthy HIV-1 uninfected, viremic untreated and long-term treated HIV-1 infected individuals and their relationship with PD-1^+^ CD4 T cells. The frequency of CD32^+^ CD4 T cells was increased in viremic as compared to treated individuals in LNs and a large proportion(up to 50%) of CD32^+^ cells co-expressed PD-1 and were enriched within T follicular helper cells(Tfh) cells. We next investigated the role of LN CD32^+^ CD4 T cells in the HIV reservoir. Total HIV DNA was enriched in CD32^+^ and PD-1^+^ CD4 T cells as compared to CD32^-^ and PD-1^-^ cells in both viremic and treated individuals but there was no difference between CD32^+^ and PD-1^+^ cells. There was not enrichment of latently infected cells with inducible HIV-1 in CD32^+^ versus PD-1^+^ cells in ART treated individuals. HIV-1 transcription was then analyzed in LN memory CD4 T cell populations sorted on the basis of CD32 and PD-1 expression. CD32^+^PD-1^+^ CD4 T cells were significantly enriched in cell associated HIV RNA as compared to CD32^-^PD-1^-(^average 5.2 fold in treated and 86.6 fold in viremics), to CD32^+^PD-1^-(^2.2 fold in treated and 4.3 fold in viremics) and to CD32^-^PD-1^+^ cell populations(2.2 fold in ART treated and 4.6 fold in viremics). Similar levels of HIV-1 transcription were found in CD32^+^PD-1^-^ and CD32^-^PD-1^+^ CD4 T cells. Interestingly, the proportion of CD32^+^ and PD-1^+^ CD4 T cells negatively correlated with CD4 T cell counts and length of therapy while positively correlated with viremia. Therefore, the expression of CD32 identifies, independently of PD-1, a CD4 T cell population with persistent HIV-1 transcription and CD32 and PD-1 co-expression the CD4 T cell population with the highest levels of HIV-1 transcription in both viremic and treated individuals.

**Importance:** The existence of long-lived latently infected resting memory CD4 T cells represents a major obstacle to the eradication of HIV infection. Identifying cell markers defining latently infected cells containing replication competent virus is important in order to determine the mechanisms of HIV persistence and to develop novel therapeutic strategies to cure HIV infection. We provide evidence that PD-1 and CD32 may have a complementary role in better defining CD4 T cell populations infected with HIV-1. Furthermore, CD4 T cells co-expressing CD32 and PD-1 identify a CD4 T cell population with high levels of persistent HIV-1 transcription.

## INTRODUCTION

The existence of long-lived latently HIV-1 infected resting memory CD4 T cells represents a major obstacle to the eradication of HIV infection(1-6). Several studies in the recent years have greatly contributed to the characterization of different CD4 T cell populations containing inducible replication competent HIV-1. In the blood, central memory CD4 T cells defined by the CD45RA^-^ CCR7^+^CD27^+^ phenotype and transitional memory CD4 T cells defined by CD45RA^-^CCR7^-^CD27^+^ phenotype have been identified as major cellular compartments of the latent HIV-1 reservoir while CD4 T cells with stem-cell like properties as a minor component(7, 8). Recent studies always performed in blood has also shown that a series of markers such as HLA-DR, CCR6, TIGIT and CD30 help to define CD4 T cell populations infected with HIV-1(9-11). Studies performed in memory CD4 T cell populations isolated from LNs of viremic and long-term treated HIV-1 infected individuals have demonstrated that PD-1 positive and Tfh CD4 T cells are the major reservoir for replication competent and infectious virus in both viremic and long-term treated individuals(12, 13). A recent study performed in blood has proposed that the low affinity receptor for the immunoglobulin G Fc fragment, CD32a, defines a small population of HIV-1 infected resting CD4 T cells and therefore CD32a^+^ CD4 T cells represent the elusive HIV reservoir(14). A recent study has challenged the findings that CD32a is a marker defining the elusive HIV reservoir(15). CD32^+^ CD4 T cells expressed several markers of activation, they are not enriched in HIV DNA while they are enriched for transcriptionally active HIV in long-term treated individuals. Along the same line, another study performed in blood has also failed to show enrichment of HIV infected CD4 T cells within CD32^+^ cells(16).

On the basis of the CD32a study and of our previous observations on the role of PD-1^+^/Tfh cells in the HIV reservoir(12-14), we have addressed the following issues: a) the differences in the percentage of CD32^+^ CD4 T cells in healthy versus HIV infected individuals, b) the distribution of CD32 in blood versus lymphoid tissue, i.e. lymph nodes(LNs), c) the distribution of CD32 in different populations of memory CD4 T cells, and d) the relationship between CD32^+^ and PD-1^+^ CD4 T cell populations and their role in defining the HIV reservoir.

## RESULTS

### Distribution of CD32+ CD4 T cells in blood and lymph nodes

Blood and LN biopsies were obtained from 9 healthy subjects, 19 long-term antiretroviral therapy(ART) treated and 10 viremic individuals (Table 1). In healthy individuals, lymph node biopsies were obtained from patients undergoing vascular surgery. A small percentage (<1%) of memory CD4 T cells in blood expressed CD32 but there were not significant differences between healthy, long-term ART treated and viremic individuals (Fig 1A and B). Memory CD4 T cells from LNs of the same individuals contained significantly greater percentages of CD32^+^ cells as compared to blood (healthy *P* = 0.003, ART treated *P* <0.0001, viremics *P* = 0.002). The percentage of LNs CD32^+^ CD4 T cells was significantly greater in viremics versus long-term ART treated individuals (Fig 1A and (B) (*P* = 0.0003) but not versus healthy individuals (Fig 1A and B).

**Table 1.** Study cohort: clinical data.

| Patient ID | Age | Sex | Duration of HIV infection (years) | Years of suppressive ART | Viral load (copies/ml) | CD4 count (cells/μl) | ART Status |
| --- | --- | --- | --- | --- | --- | --- | --- |
| MP106 | 37 | M | 0.1 | 0 | 160'000 | 501 | Treatment Naive |
| MP140 | 23 | M | 0.08 | 0 | 360'000 | 427 | Treatment Naive |
| MP119 | 23 | M | 1.3 | 0 | 13'000 | 498 | Treatment Naive |
| MP124 | 46 | F | 0.06 | 0 | 510'000 | 468 | Treatment Naive |
| MP113 | 33 | M | 31.98 | 0 | 640 | 594 | Treatment Naive |
| MP117 | 51 | M | 5.42 | 0 | 54'000 | 504 | Treatment Naive |
| MP125 | 35 | M | 0.16 | 0 | 17'000 | 511 | Treatment Naive |
| MP118 | 29 | M | 0.06 | 0 | 14'000 | 538 | Treatment Naive |
| SA150 | 53 | M | 10 | 0 | 6'900 | 578 | Treatment Naive |
| SA143 | 37 | M | 0.3 | 0 | 25'000 | 704 | Treatment Naive |
| MP129 | 44 | M | 14.85 | 14.71 | <20 | 928 | ART Treated |
| MP068-1 | 45 | M | 3.33 | 3 | <20 | 928 | ART Treated |
| MP136 | 44 | M | 7.5 | 6 | <20 | 732 | ART Treated |
| MP123 | 61 | M | 9.25 | 9 | <20 | 1270 | ART Treated |
| MP137 | 39 | M | 3.59 | 2.84 | 24 | 700 | ART Treated |
| MP064 | 36 | M | 7.88 | 4.49 | <20 | 719 | ART Treated |
| MP141 | 31 | M | 11.56 | 1.5 | <20 | 558 | ART Treated |
| MP094 | 47 | M | 8 | 8 | <20 | 451 | ART Treated |
| MP058 | 46 | F | 10.86 | 10 | <20 | 666 | ART Treated |
| MP082 | 53 | M | 22.45 | 8 | <20 | 1236 | ART Treated |
| MP096 | 54 | F | 18 | 13 | <20 | 1253 | ART Treated |
| MP103 | 41 | M | 1.62 | 1.16 | <20 | 217 | ART Treated |
| MP068-2 | 47 | M | 5.12 | 5 | <20 | 487 | ART Treated |
| MP060 | 42 | M | 21 | 8 | <20 | 786 | ART Treated |
| MP067 | 41 | M | 14.01 | 14 | <20 | 609 | ART Treated |
| MP072 | 47 | M | 18 | 2 | <20 | 614 | ART Treated |
| MP070 | 55 | M | 7.06 | 2 | <20 | 615 | ART Treated |
| MP100 | 36 | F | 13.06 | 10 | <20 | 643 | ART Treated |
| MP145 | 39 | M | 8.35 | 4 | <20 | 820 | ART Treated |

**Fig 1.**
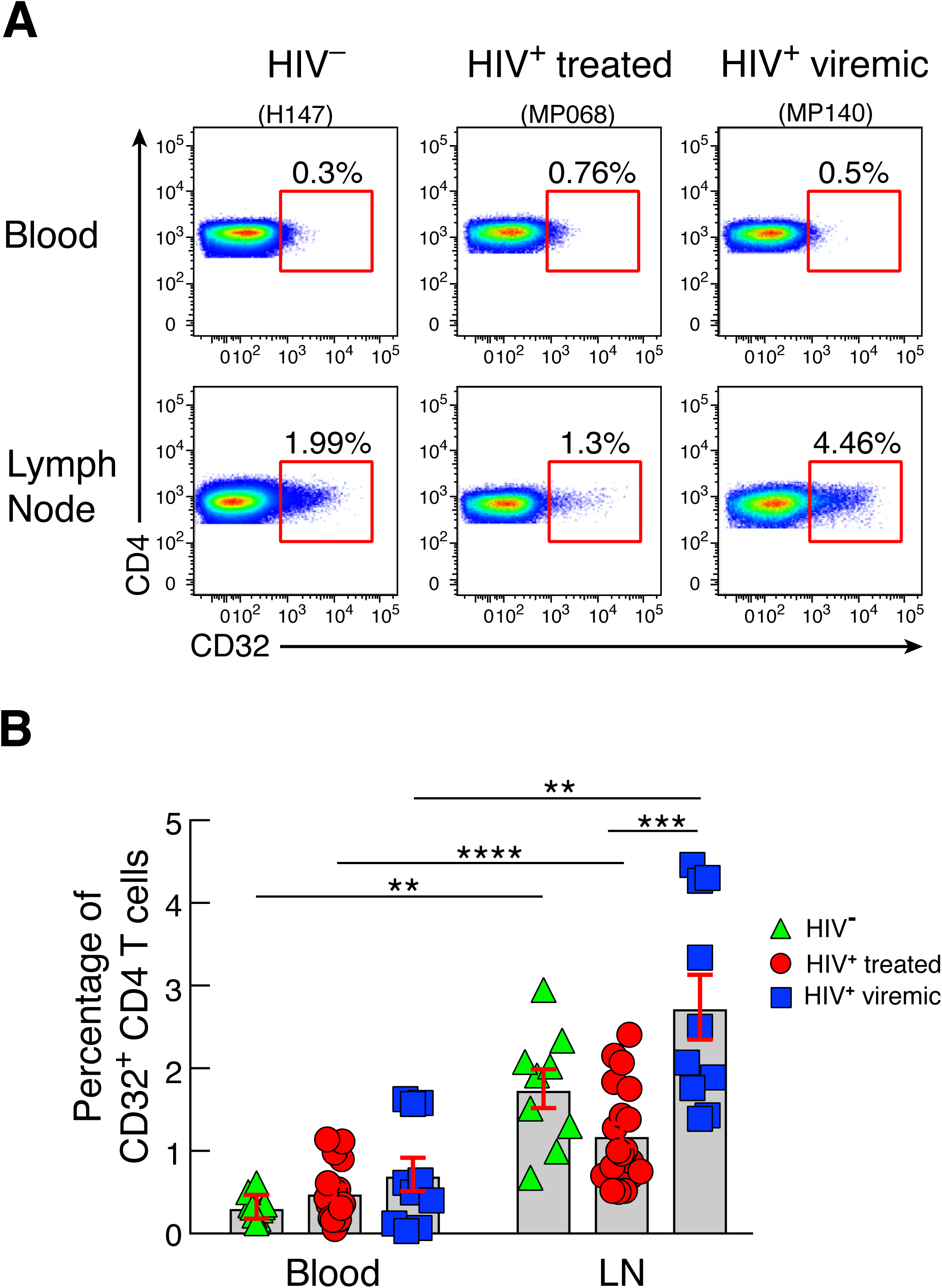
CD32 expression on CD4 T cells in LN and blood of HIV-1 infected and uninfected individuals. LN and PB mononuclear cells were isolated from the same HIV-uninfected (N=9), HIV-1 infected ART treated (N=19) and HIV-1 infected viremic individuals (N=10). Cells were stained with anti-CD3, anti-CD4, anti-CD45RA, anti-CD32 antibodies.(**A**) Representative flow cytometry profiles of blood and LN memory (CD45RA^-^) CD4 T cell populations expressing CD32 in representative HIV-1 uninfected, ART treated and viremic subjects.(**B**) Cumulative data on the frequencies of CD32^+^ memory CD4 T cells in blood and LN mononuclear cells of HIV-1 uninfected (green), ART treated (red) and viremic (blue) individuals. *P* values were obtained by Mann-Whitney to compare the three groups and Wilcoxon signed-rank test to compare frequencies between blood and LNs. \*\**P*<0.01, *** *P*<0.001, \*\*\*\**P*<0.0001. Error bars denote mean ± S.E.M.

These results indicate that CD32 expression in memory CD4 T cells from blood and LNs is not restricted to HIV-1 infected individuals and that CD32^+^ CD4 T cells are greatly enriched in LNs.

### Distribution of CD32 in LN and blood memory CD4 T cell populations

Next, we investigated the distribution of CD32 in different memory LN CD4 T cell populations defined by the expression of CXCR5 and PD-1 in healthy, long-term ART treated and viremic individuals. Memory CD32^+^ CD4 T cells were consistently enriched in PD-1^+^CXCR5^-^ and PD-1^+^CXCR5^+^ CD4 T cell populations in the three study groups (P <0.05 for all study groups) (Fig 2A and B). The PD-1^+^CXCR5^+^ CD4 T cells correspond to Tfh cells (12, 17-19). PD-1^+^ CD4 T cells were greatly enriched within the CD32^+^ CD4 T cell population in the three study groups and 40 and 50% of PD-1^+^ CD4 T cells co-expressed PD-1 and CD32 in long-term ART treated and viremic individuals, respectively (Fig 2C) (*P* <0.0001 in healthy and ART treated and *P* = 0.002 in viremics). Similarly, a substantial proportion of Tfh cells ranging between 10% in ART treated and 20% in viremic individuals were contained within the CD32^+^ CD4 T cell population (Fig 2D) (*P* <0.0001 and *P* = 0.002, respectively). However, as compared to the CD32^+^ CD4 T cell population, Tfh cells were significantly enriched within the PD-1^+^ CD4 T cell population (*P* <0.0001) (S1 Fig). The distribution of memory CD4 T cell populations defined by CD32 and PD-1 was also investigated in blood (S2 Fig). The CD32^-^PD-1^-^ CD4 T cell population was significantly reduced in viremic individuals as compared to the healthy subjects (*P* = 0.0007). No differences in the percentage of CD32^+^PD-1^-^ and CD32^+^PD-1^+^ CD4 T cells were observed in the three study groups while CD32^-^PD-1^+^ CD4 T cells significantly increased in viremic individuals (13.1% in viremics versus 4% in healthy, *P* = 0.0007 and 6.1% in treated, *P* = 0.006).

**Fig 2.**
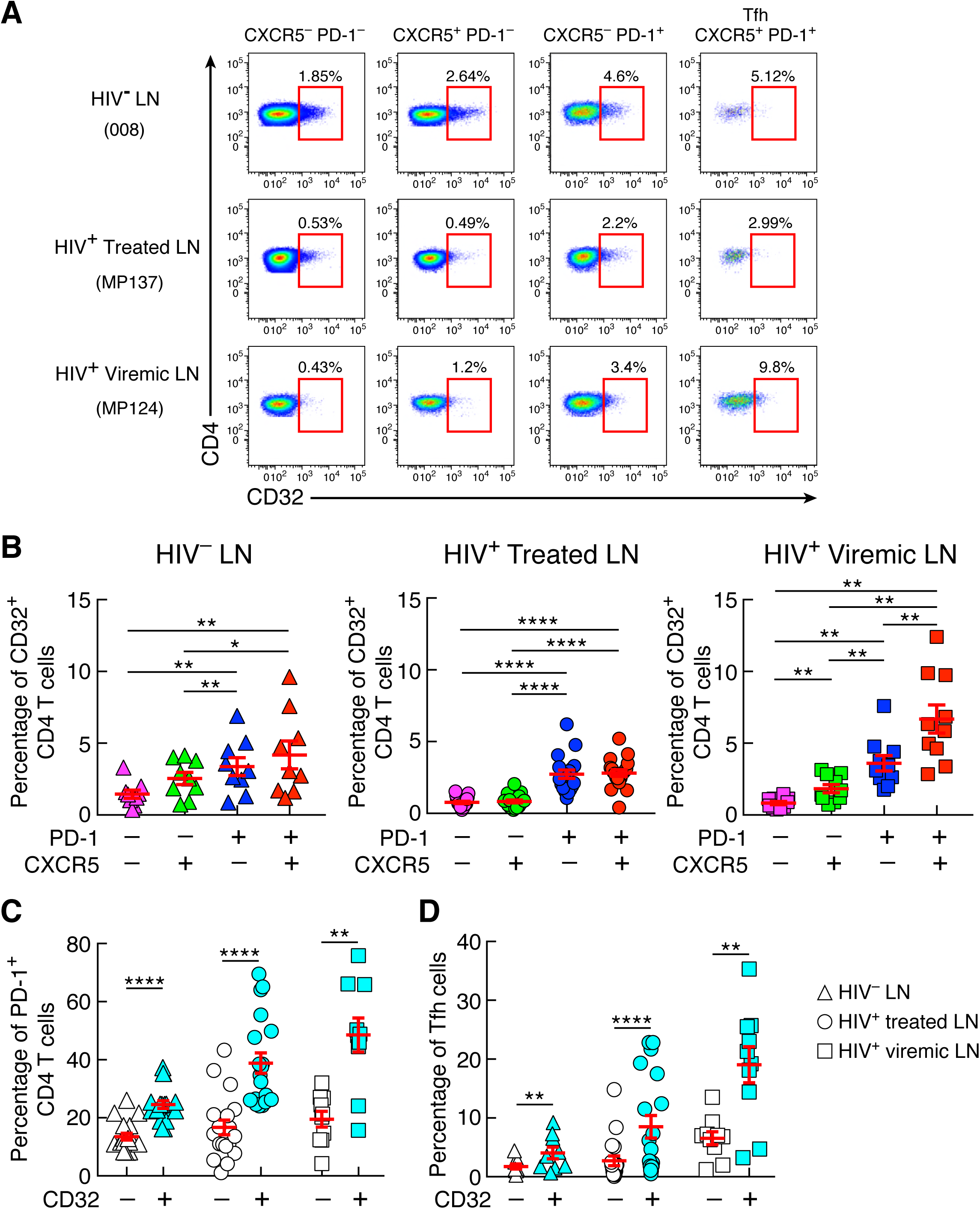
CD32+ CD4 T cells from LNs are enriched in PD-1+ and Tfh cells. (**A**) Representative flow cytometry profiles of CD32 expression in LN memory CD4 T cell populations defined by PD-1 and CXCR5 expression of HIV-1 uninfected, HIV-1 infected ART treated and viremic individuals.(**B**) Cumulative data on the frequencies of CD32^+^ memory CD4 T cells within PD-1^-^ CXCR5^-^, PD-1^-^CXCR5^+^, PD-1^+^CXCR5^-^ and Tfh (PD-1^+^ CXCR5^+^) cell populations isolated from LNs of 9 HIV-1 uninfected, 19 ART treated and 10 viremic individuals.(**C**) Percentage of PD-1^+^ cells in gated CD32^-^ and CD32^+^ memory (CD45RA^-^) CD4 T cell populations in the three groups. (**D**) Distribution of Tfh within CD32^-^ and CD32^+^ memory CD4 T cell populations. Triangles represent HIV-1 uninfected, round circles represent ART treated and squares represent viremic individuals. \**P*<0.05, \*\**P*<0.01, \*\*\*\**P*<0.0001 values were obtained by Wilcoxon signed-rank test and error bars denote mean ± S.E.M.

### Relationship between CD32+ and PD-1+ CD4 T cell populations

The simultaneous assessment by mass cytometry of 30 phenotypic markers defining different cell lineages, memory differentiation, cell activation and cell trafficking has provided the possibility to analyzed more in depth the differences between CD32^+^ and CD32^-^ lymph node memory CD4 T cell populations in the three study groups. The large majority (13 out of 16) of markers defining chemokine receptors (CXCR3, CCR6, CCR4), cell activation (CD25, CD38, HLA-DR), memory cell differentiation (ICOS, CD57, CD27, CD127, CXCR5, PD-1, CD40L) and HIV-1 co-receptors (CCR5 and CXCR4) were greatly increased in CD32^+^ as compared to CD32^-^ CD4 T cells regardless of the study group (Fig 3A and S3 Fig). Only CCR7, CD127 and CD27 were unchanged. The heat maps in Fig 3A show also the differences among the study groups. ICOS, CD38, PD-1 are enhanced within the CD32^+^ cells (*P* < 0.0001) while CCR4 and CD127 were slightly reduced within the CD32^-^ cells in viremic individuals. Several markers are downregulated in ART treated individuals as compared even to healthy individuals.

**Fig 3.**
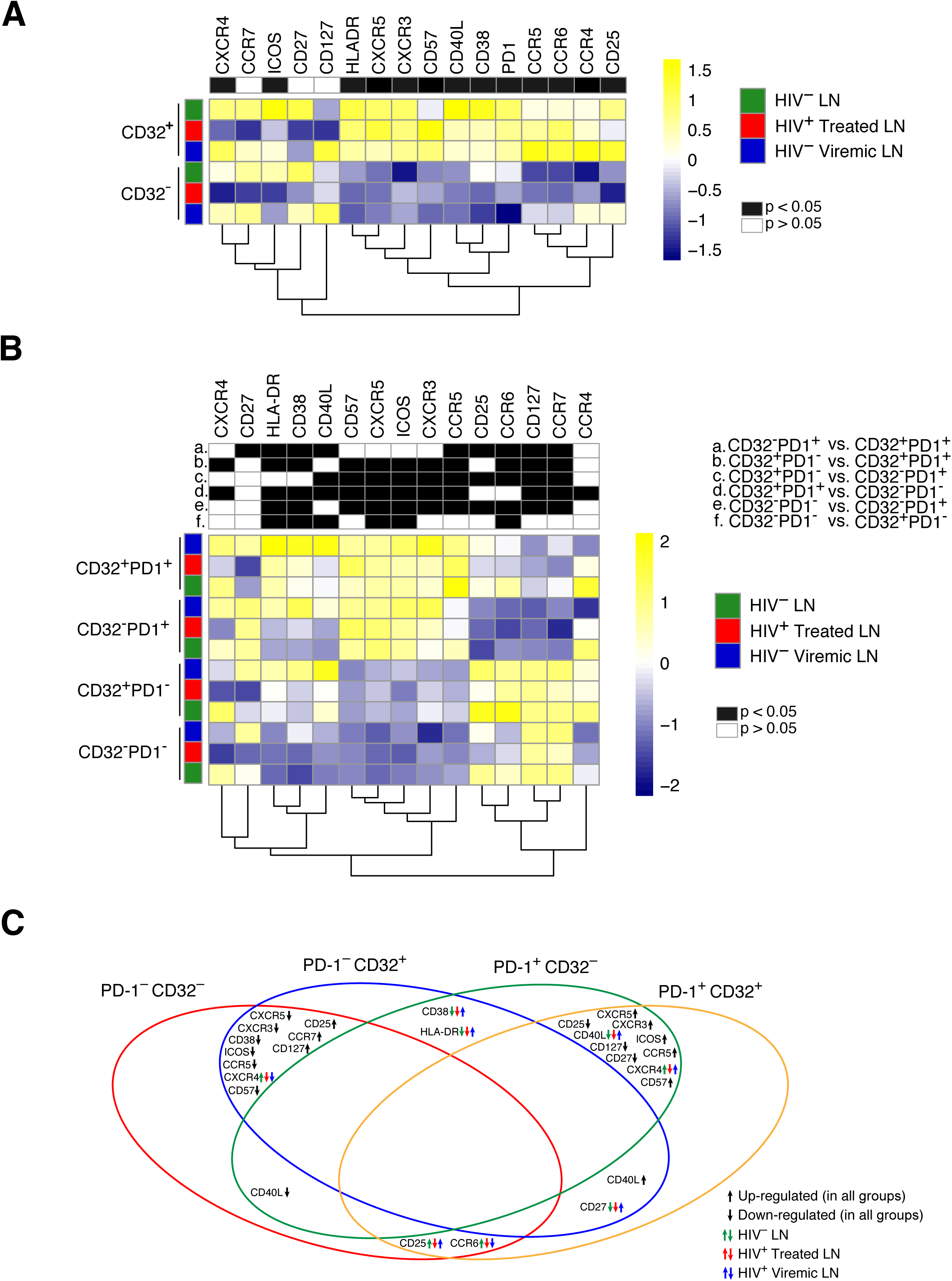
Mass cytometry analysis of LN memory CD4 T cells defined by CD32 and/or PD-1 expression. Mass cytometry staining was performed in LN mononuclear cells isolated from 9 HIV-1 uninfected, 16 HIV-1 infected ART treated and 9 viremic individuals. Cells were stained with a panel of 30 cell surface markers (S1 Table). Heat maps of mean marker intensities in memory CD4 T cells defined on the basis of (**A**) CD32 and of (**B**) CD32 and PD-1 expression.(**C**) Venn diagram summarizing common trends in marker expression between memory (CD45RA^-^) CD4 T cells defined on the basis of PD-1 and CD32 expression. Black arrows indicate trends that are common to all study groups (HIV uninfected, ART treated and viremic individuals). Green, red and blue arrows indicate trends that are specific to each study group.

Due to the overlap between CD32^+^ and PD-1^+^ CD4 T cell populations, the distribution of the same panel of makers was analyzed in lymph node memory CD4 cell sub-populations defined by the expression of CD32 and PD-1 in the three study groups (Fig 3B). The heat maps and the Venn diagram show that the expression of PD-1 determines a clear separation between the four cell populations. It appears that PD-1^-^CD32^-^ and PD-1^-^CD32^+^ CD4 T cells from one side and PD-1^+^ CD32^-^ and PD-1^+^CD32^+^ CD4 T cells from the other side share a similar phenotype and are closely related to each other (Fig 3B and C). The greater expression of markers such as CXCR5, ICOS, CD57, CD40L, CD38, HLA-DR and reduced expression of CD127 and CCR7 indicates that PD-1^+^ CD4 T cell populations are at an advanced stage of differentiation and activation. Of note, the expression of the HIV-1 co-receptors, CCR5 and CXCR4, is greatly enhanced in the PD-1^+^ CD4 T cell populations (CCR5^+^ cells 21.8% in CD32^-^PD-1^+^ and 37% in CD32^+^PD-1^+^; CXCR4^+^ ce1ls 14.2% in CD32^-^PD-1^+^ and 18% in CD32^+^PD-1^+^). Cumulative data on the expression of the 15 markers in the four memory CD4 T cell populations defined by the expression of PD-1 and CD32 independently of the study group are shown in S4 Fig.

Taken together these results indicate that there is great overlap between CD32^+^ and PD-1^+^ memory CD4 T cell populations in LNs of HIV-1 infected individuals. Importantly, the expression of PD-1 independently of CD32 expression helps to define CD4 T cells potentially more susceptible to HIV infection.

### Role of PD-1+ and CD32+ memory CD4 T cell populations in the HIV Reservoir

Having defined the distribution and the phenotypic relationship between PD-1^+^ and CD32^+^ memory CD4 T cell populations in blood and LNs, we sought to investigate the role of these cell populations in the HIV reservoir through determining whether there is: a) an enrichment of HIV DNA containing cells in LN CD32^+^ and PD-1^+^ CD4 T cells, b) a difference in the frequency of HIV DNA containing cells between LN CD32^+^ and PD-1^+^ CD4 cells in viremic and long-term ART treated individuals, c) a difference in the frequency of cells containing replication competent HIV between LN CD32^+^ and PD-1^+^ CD4 T cells in long-term treated individuals, d) a difference in HIV transcription between LN CD32^+^ and PD-1^+^ CD4 T cells in long-term ART treated individuals. Memory CD4 T cells (CD45RA-) isolated from lymph nodes of seven long-term ART treated and seven viremic individuals were sorted on the basis of CD32 and/or PD-1 expression and analyzed for the presence of total HIV DNA. A caveat of this analysis is that because of the overlapping between CD32^+^ and PD-1^+^ CD4 T cell populations, a proportion of cells co-expressing PD-1 and CD32 are present in both sorted CD32^+^ and PD-1^+^ cell populations (S5A Fig). However, because of the limited number of cells particularly in LNs of ART treated individuals, it was not possible to assess the total HIV DNA in the four cell populations defined by the expression of CD32 and PD-1(CD32^-^PD-1^-^, CD32^+^PD-1^-^, CD32^-^PD-1^+^ and CD32^+^PD-1^+^). CD32^+^ and PD-1^+^CD4 T cell populations were enriched in cells containing HIV DNA as compared to CD32^-^ and PD-1^-^ cell populations. With regard to CD32^+^ versus CD32^-^ the differences were significant in the ART treated individuals (1.83 fold, *P* = 0.01) and viremics (23.7 fold, *P* = 0.01) (Fig 4A). With regard to PD-1^+^ versus PD-1^-^ CD4 T cell populations, there was only a trend toward higher HIV DNA in PD-1^+^ versus PD-1^-^ in ART treated individuals while the differences were significant in viremic individuals (6.5 fold, *P* = 0.01) (Fig 4A).

**Fig 4.**
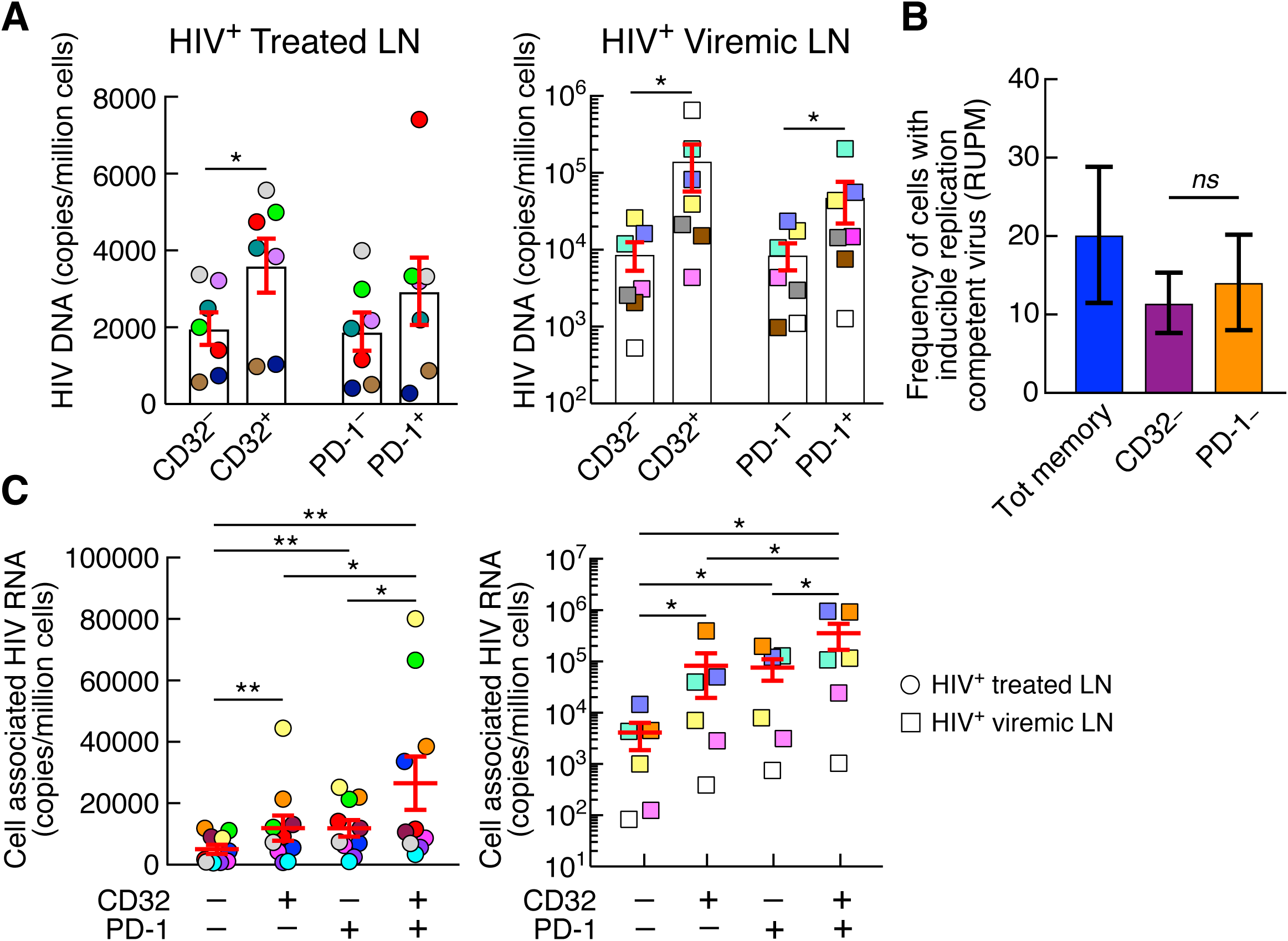
Levels of total HIV DNA and cell-associated HIV RNA in CD32+ and PD-1+ CD4 T cell populations. **(A)** Quantification of total HIV DNA(copies per million cells) of sorted memory (CD45RA^-^) CD4 T expressing or not CD32 and PD-1 from 7 aviremic ART treated (round circles) and 7 viremic individuals (squares).(**B**) Frequencies of cells with inducible replication competent HIV in sorted CD32^-^, PD-1^-^ and total memory CD4 T cells estimated using conventional limiting dilution methods by Extreme Limiting Dilution analysis (http://bioinf.wehi.edu.au/software/elda/) (32, 33) in 3 aviremic ART treated individuals.(**C**) Levels of cell-associated unspliced HIV RNA(copies/million cells) in sorted CD32^-^PD-1^-^, CD32^+^PD-1^-^, CD32^-^PD-1^+^ and CD32^+^PD-1^+^ memory CD4 T cell populations isolated from 10 aviremic ART treated and 6 viremic individuals. \**P*<0.05, \*\**P*<0.01 values were obtained by Wilcoxon signed-rank test and error bars denote mean ± S.E.M.

These results show an enrichment of HIV infected cells within CD32^+^ and PD-1^+^ LN memory CD4 T cells but not substantial difference between these cell populations. The massive proportion of cells infected on the basis of the total HIV DNA content in CD32^+^ and PD-1^+^ CD4 T cells are consistent with the phenotypic data (Fig 3) showing increased expression of HIV-1 co-receptors in these cell populations and indicate that they may serve as a major reservoir of HIV-1.

To estimate the frequencies of HIV-1 infected cells containing inducible replication competent virus, a quantitative virus outgrowth assay (QVOA) was performed in three ART treated individuals. The objective of the QVOA was to determine whether the exclusion of CD32^+^ or PD-1^+^ cells had a different impact on the frequency of HIV-1 infected cells with inducible virus. Because of the limited number of cells isolated from ART treated individuals, it was not possible to perform the QVOA on CD32^+^ and PD-1^+^ CD4 T cell populations. For the QVOA, different cell concentrations (five-fold limiting dilutions: 10^5^, 2.10^4^ and 4.10^3^) and multiple replicates of sorted viable lymph node CD32^-^, PD-1^-^ and total memory (CD45RA^-^) CD4 T cells (S5A Fig) were co-cultured with allogeneic fresh CD8 T cell-depleted blood mononuclear cells from HIV-1 uninfected individuals and frequencies were estimated at day 14 using Extreme Limiting Dilution Assay (20). These analyses provide the average RNA-unit per million (RUPM) for each cell population investigated. The frequencies of cells with inducible HIV RNA in CD32^-^ and PD-1^-^ CD4 cell populations were not significantly different (Fig 4B). In order to interpret correctly these results, it is important to consider that sorting out CD32^+^ cells there are still PD-1^+^ cells remaining in the sorted CD32-cell population (S5A Fig). Since also PD-1^+^ cells contain inducible HIV, it is expected the finding that there is a trend towards reduction of the frequency of cells with inducible HIV and not a more dramatic reduction. Along the same line, by sorting out PD-1^+^ cells there are still CD32^+^ cells remaining in the sorted PD-1^-^ cell population which explains the partial reduction found. Finally, there was a trend towards higher frequencies of cells with inducible HIV RNA in the total memory CD4 T cell population as compared to CD32^-^ and PD-1^-^ populations (Fig 4B).

We have recently shown that PD-1^+^ CD4 T cells serve as the major CD4 T cell compartment in blood and LNs for replication competent and infectious HIV-1 and for active and persistent virus transcription in long-term ART treated aviremic individuals (13). We then investigated whether there was a difference in HIV transcription between LN CD32^+^ and PD-1^+^ CD4 T cells in ten long-term ART treated and six viremic individuals. To determine the cell compartment (s) serving as sites of active and persistent HIV-1 transcription, cell-associated HIV-1 RNA was assessed in LN memory CD32^-^PD-1^-^, CD32^+^PD-1^-^, CD32^-^PD-1^+^ and CD32^+^PD-1^+^ CD4 T cell populations. The gating strategy for cell sorting is shown in S5B. The results indicate that the levels of cell-associated HIV-1 RNA were significantly higher in CD32^+^PD-1^+^ CD4 T cells as compared to the other three cell populations (Fig 4C). CD32^+^PD-1^+^ CD4 T cells were enriched in cell-associated HIV-1 RNA as compared to CD32^-^PD-1^-(^average 5.2 fold in ART treated and 86.6 fold in viremics), to CD32^+^PD-1^-(^2.2 fold in ART treated and 4.3 fold in viremics) and to CD32^-^PD-1^+^ cell populations (average 2.2 fold in ART treated and 4.6 fold in viremics). CD32^+^PD-1^-^ and CD32^-^ PD-1^+^ cell populations had also significantly higher levels of cell-associated HIV RNA as compared to CD32^-^PD-1^-^ cells (*P* < 0.05) while no differences were found between CD32^+^PD-1^-^ and CD32^-^PD-1^+^ cell populations.

Therefore, the co-expression of CD32 and PD-1 identifies a population with greater HIV transcriptional activity within Tfh cells and the sole expression of CD32 identifies a cell population with levels of transcriptional activity similar to those measured in single PD-1^+^ CD4 T cells.

Recent results (21) have suggested that the expression of CD32 on blood CD4^+^ T cells was reflective of either the presence of doublets or partial exchange of a B cell and CD4^+^ T cell membranes. We also have carefully addressed this issue in our flow cytometry experiments. The exclusion of doublets is a mandatory step in the flow cytometry analysis of cell populations. As shown in S6 Fig we have totally excluded that the expression of CD32 on lymph node CD4^+^ T cells was the result of B-T cells doublets. In the presence of B-CD4 T cell doublets we should find that the expression of CD32 in the different memory CD4 T cell populations would also be associated with the expression of CD19. The doublets have been excluded in two sequential analyses. Similarly, CD19^+^ B cells and CD8^+^ T cells are excluded when gating for CD4^+^ T cells (S6A Fig). S6B Fig clearly demonstrates that there is not CD19 expression on single CD32^+^ CD4 T cells as well as CD32^+^PD-1^+^ CD4 T cells.

### CD32+ and PD-1+ CD4 T cells and markers of HIV disease activity

We then determined the relationship between CD32^+^ and PD-1^+^ CD4 T cells and indicators of HIV disease activity and the length of suppressive ART. The percentage of CD32^+^ CD4 T cells negatively correlated with CD4 T cell count (*r* = −0.58, *P* = 0.0008) and length of ART treatment (*r* = −0.79, *P* <0.0001) while positively correlated with the levels of viremia (*r* = 0.68, *P* <0.0001) (Fig 5A). Next, we analyzed the relationships between the levels of viremia and the length of treatment and the percentage of the four LN memory CD4 T cell populations sorted on the basis of the expression of CD32 and PD-1. Interestingly, the percentage of CD32^-^PD-1^-^ cells negatively correlated with the levels of viremia (*r* = −0.46, *P* = 0.01) while the percentages of CD32^+^PD-1^-^, CD32^-^PD-1^+^ and CD32^+^PD-1^+^ cells strongly correlated with viremia (*r* = 0.56, *P* = 0.001; *r* = 0.46, *P* = 0.03; *r* = 0.65, *P* = 0.0001, respectively) (Fig 5B). In contrast, the percentage of CD32^-^PD-1^-^ cells positively correlated with the length of ART(*r* = 0.6, *P* = 0.0004) while the percentage of the other three cell populations negatively correlated with the length of ART treatment (*r* = −0.67, *P* <0.0001; *r* = −0.56, *P* = 0.001; *r* = −0.8, *P*<0.0001, respectively) (Fig 5C).

**Fig 5.**
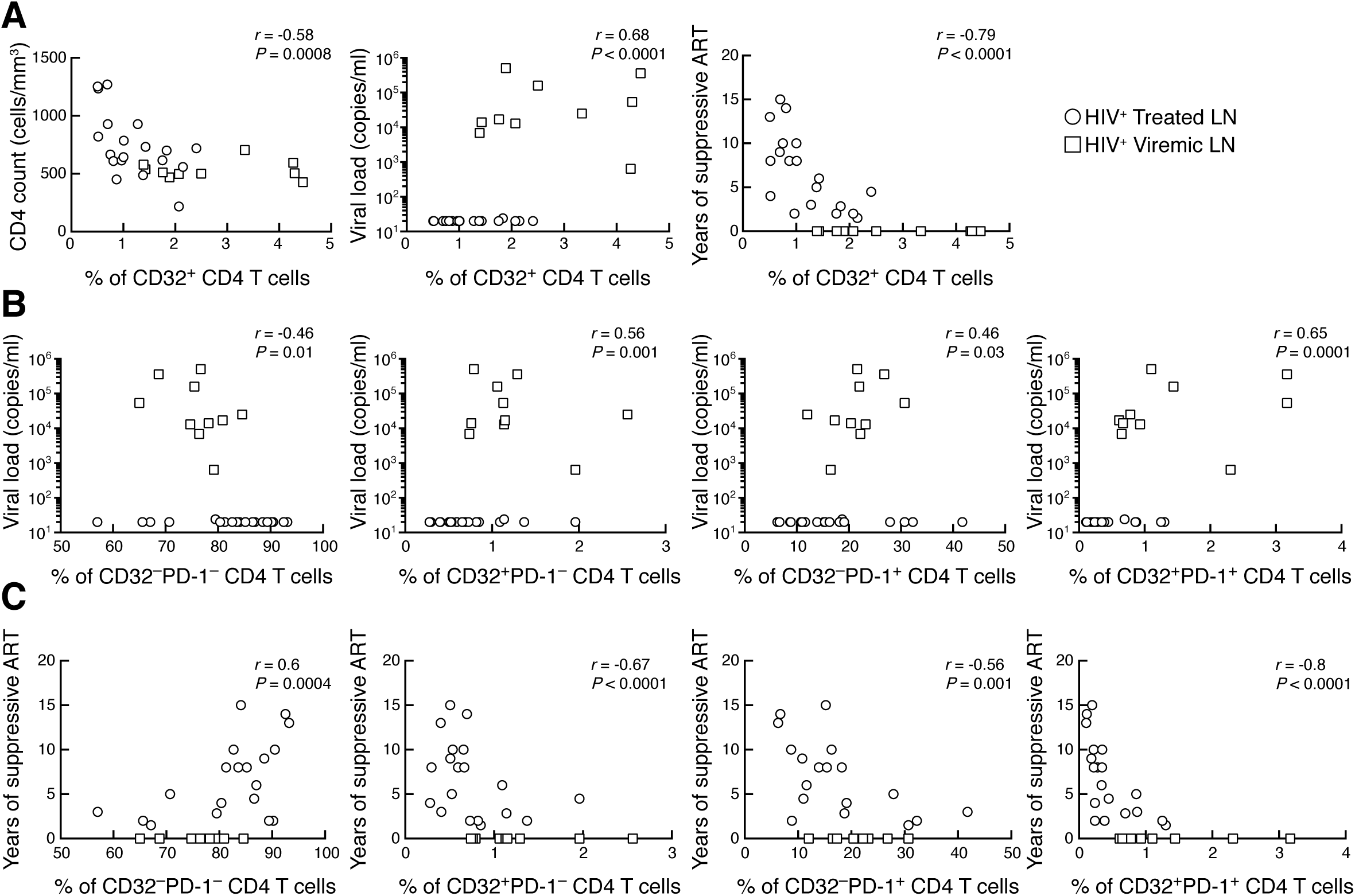
Correlations between memory CD4 T cell populations defined by CD32 and/or PD-1 and parameters of disease activity. (**A**) Correlation between the percentage of CD32^+^ memory CD4 T cells and CD4 T cell count, levels of viremia and years of suppressive ART. Correlation between the percentage of CD32^-^PD-1^-^, CD32^+^PD-1^-^, CD32^-^PD-1^+^ and CD32^+^PD-1^+^ populations and (**B**) levels of viremia (**C**) and years of suppressive ART. Spearman rank test was used for correlations.

A similar pattern was observed for PD-1^+^ CD4 T cells (Fig 6). The percentage of PD-1^+^ CD4 T cells negatively correlated with CD4 T cells count (*r* = −0.51, *P* = 0.004) and length of suppressive ART(*r* = −0.58, *P* = 0.0009) while positively correlated with the levels of viremia (*r* = 0.43, *P* = 0.01).

**Fig 6.**
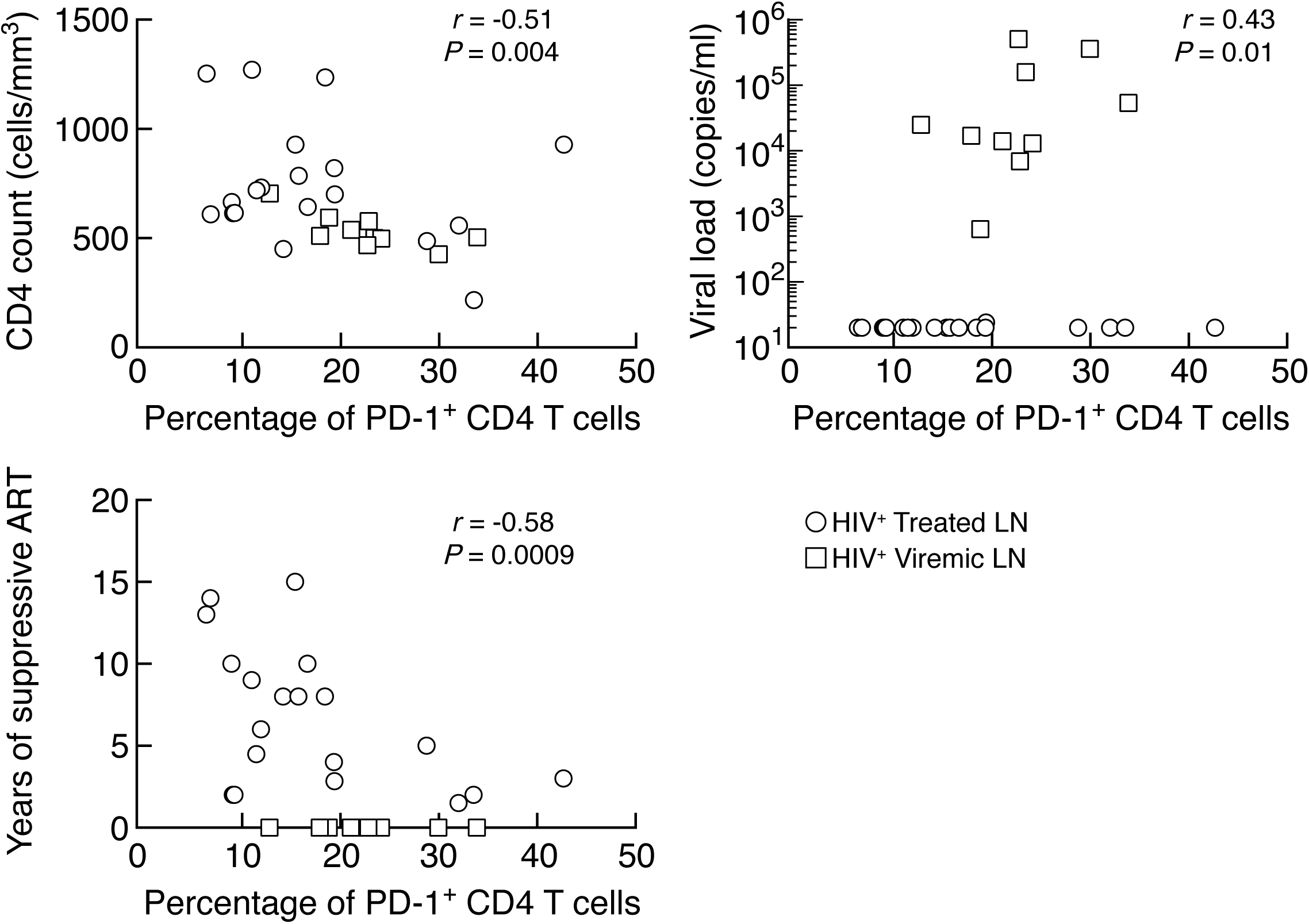
Correlation between memory CD4 T cell populations defined by PD-1 and parameters of disease activity. Correlation between the percentage of PD-1^+^ memory CD4 T cells with CD4 T cell count, viral load and years of suppressive ART. Spearman rank test was used for correlations.

These results indicate that the expansion of both CD32^+^ and PD-1^+^ memory CD4 T cell populations in lymph nodes is driven by HIV replication.

## DISCUSSION

The percentage (range of 0.2-0.5%) of CD32^+^ CD4 T cells and its intensity of expression are extremely low in blood thus rendering challenging to perform in depth characterization of circulating CD32^+^ CD4 T cells. For these reasons, we decided to study CD32^+^ CD4 T cells in LNs. Lymphoid tissues are the major anatomic compartment for HIV replication, spreading and persistence and therefore they are the ideal anatomic compartment to understand the role of CD32^+^ CD4 T cells in the HIV reservoir and their relationship with PD-1^+^/Tfh cells that have been shown to serve as a major reservoir for HIV(13, 22-29). The percentages of CD32^+^ CD4 T cells were similar in healthy HIV uninfected, viremic and ART treated individuals in the blood. Higher percentage of CD32^+^ CD4 T cells were found in LNs in the three study groups and were significantly expanded in viremic individuals but similar percentages were found in LNs of healthy and treated individuals thus indicating that CD32 expression on CD4 T cells is not exclusive of HIV infection as proposed by Descours et al (14).

CD32^+^ and PD-1^+^ CD4 T cell populations are tightly correlated as indicated by the large proportion (up to 50%) of CD32^+^ CD4 T cells co-expressing PD-1. The co-expression of PD-1 is important for understanding the status of activation and differentiation of CD32^+^ cells and their role in the HIV reservoir. Indeed, the expression of PD-1 together with HLA-DR and CD38 in CD32^-^PD-1^+^ and CD32^+^PD-1^+^ CD4 T cells indicates that these cell populations are associated with high levels of cell activation and advanced differentiation as compared to the CD32^-^PD-1^-^ and CD32^+^PD-1^-^cell populations. Of note, CD32^+^ and PD-1^+^ CD4 T cells expressed higher levels of the HIV-1 co-receptors CCR5 and CXCR4 thus rendering these cells a preferential target for HIV infection. These data are consistent with two recent studies (15, 16) showing that CD32^+^ CD4 T cells express higher levels of activation markers and HIV co-receptors.

On the basis of the quantification of the total HIV DNA and of cells with inducible replication competent HIV, CD32^+^ and PD-1^+^ CD4 T cells contribute equally to the HIV reservoir in long-term treated individuals and no differences were also found in viremic individuals. More importantly, the analysis of cell-associated HIV RNA has demonstrated equal levels of persistent HIV transcription in CD32^+^PD-1^-^ and CD32^-^PD-1^+^ CD4 T cell populations and significantly higher levels of transcription in the CD32^+^PD-1^+^ cells. These findings are in agreement with a recent study (15) failing to confirm the conclusions of Descours et al. that CD32 defines the latent ‘elusive’ HIV reservoir. However, CD32, independently of PD-1, defines an additional population of memory CD4 T cells with persistent HIV transcription. When co-expressed with PD-1, CD32 defines the CD4 T cell population with the highest levels of transcription.

In conclusion, the combined use of CD32 and PD-1 identify three cell populations, CD32^+^PD-1^-^, CD32^-^PD-1^+^ and CD32^+^PD-1^+^, of memory CD4 T cells responsible for most of persistent HIV transcription in long-term treated individuals.

## METERIAL AND METHODS

### Study groups

Samples from 9 healthy subjects, 19 subjects on suppressive ART and from 10 viremic individuals not receiving ART at the time of sampling were used in this study. None of the participants under ART had detectable plasma viremia at the time of the study, as assessed by viral load measurement using the Ampliprep/Cobas Taqman HIV-1 Test v 2.0(Roche), with a detection limit of 20 HIV RNA copies/ml of plasma. All participants underwent lymph node biopsy (inguinal lymph nodes) and blood was collected at the same time. Subject characteristics are summarized in Table 1. These studies were approved by the Institutional Review Board of the Centre Hospitalier Universitaire Vaudois, and all subjects gave written informed consent.

### Isolation of lymph node cells

Lymph node mononuclear cells were isolated by mechanical disruption (22) and cells were cryopreserved in liquid nitrogen.

### Antibodies

The following antibodies were used: APC-H7-conjugated anti-CD3(clone SK7, BD), PB-conjugated anti-CD19(clone HIB19, Biolegend), PB-conjugated CD8(clone RPA-T8, BD) FITC conjugated anti-CXCR5(clone RF8B2, BD), PE-Cy7-conjugated anti-PD-1(clone EH12.1, BD), Alexa700 anti-CD4(clone RPA-T4, Biolegend), APC-conjugated anti-CD32(FUN-2, Biolegend), ECD-conjugated anti-CD45RA(clone 2H4, Beckman Coulter). Aqua LIVE/DEAD stain kit was used to determine the viability of cells. Cells were acquired on an LSRII flow cytometer using the FACSDiva software (BD) and analyzed using FlowJo 9.7(TreeStar Inc) or sorted using FACSAria (BD). The CD32 antibody used in the present study is the same used in the study by Descours et al. and this antibody does not discriminate between CD32a and CD32b.

### CyTOF marker labeling and detection

Cryopreserved lymph node mononuclear cells were thawed and resuspended in complete RPMI medium (Gibco; Life Technologies; 10% heat-inactivated FBS(Institut de Biotechnologies Jacques Boy), 100 IU/ml penicillin, and 100 µg/ml streptomycin (BioConcept)) at 1’10^6^ cells/ml.

Viability of cells in 500 µl of PBS was assessed by incubation with 50 µM of cisplatin (Sigma-Aldrich) for 5 min at room temperature (RT) and quenched with 500 µl of fetal bovine serum. Next, cells were incubated for 20 minutes at 4°C with anti-CD32 APC antibody. Next, cells were washed and incubated for 30 min at RT with a 50 µl cocktail of cell surface metal conjugated antibodies (S1 Table). Cells were washed and fixed for 10 min at RT with 2.4% PFA. Total cells were identified by DNA intercalation (1 µM Cell-ID Intercalator, Fluidigm/DVS Science) in 2% PFA at 4 °C overnight. Labeled samples were assessed by the CyTOF1 instrument that was upgraded to CyTOF2(Fluidigm) using a flow rate of 0.045 ml/min. FCS files were normalized to the EQ Four Element Calibration Beads using the CyTOF software. For conventional cytometric analysis of memory CD4 T cell populations, FCS files were imported into Cytobank Data Analysis Software.

### Quantification of total HIV DNA

CD32^+^, CD32^-^, PD-1^+^ and PD-1^-^ memory (CD45RA^-^) CD4 T cell populations were sorted using FACSAria and lysates were directly used in a nested PCR to quantify both total HIV DNA and CD3 gene copy numbers, as previously described (7).

### Quantification of cell-associated RNA

Cell-associated HIV-1 RNA from individual samples was extracted from CD4 T cell populations sorted on the basis of CD32 and PD-1 expression (CD32^-^PD-1^-^, CD32^+^PD-1^-^, CD32^-^PD-1^+^, CD32^+^PD-1^+^) and subjected to DNase treatment (RNAqueous-4PCR Kit, Ambion). RNA standard curves were generated after isolation and quantification of viral RNA from supernatant of ACH2 culture as previously described (30). One-step cDNA synthesis and pre-amplification were performed as previously described (31).

### Viral outgrowth assay (VOA)

Different cell concentrations (five-fold limiting dilutions: 10^5^, 2.10^4^ and 4.10^3^ for lymph node CD4 T cells) and multiple replicates of sorted viable LN total memory (CD45RA-), CD32^-^ and PD-1^-^ memory CD4 T cells isolated from three ART treated HIV-1 infected individuals were cultured with allogenic fresh CD8-depleted blood mononuclear cells (10^6^ cells/mL) from HIV-1 uninfected subjects in the presence of anti-CD3/anti-CD28 MAb coated plates (10 µg/mL) for 3 days. Cells were carefully transferred to new uncoated plates post 3 days of activation. All cell conditions were cultured in complete RPMI for 14 days. Medium was replaced at day 5. Supernatants were collected at day 14. The presence of HIV-1 RNA was assessed by COBAS^®^ AmpliPrep/TaqMan^®^ HIV-1 Test (Roche; Switzerland). Wells with detectable HIV-1 RNA(≥20 HIV-1 RNA copies/ml) were referred to as HIV-1 RNA-positive wells. RUPM frequencies were estimated by conventional limiting dilution methods using Extreme Limiting Dilution analysis (http://bioinf.wehi.edu.au/software/elda/) (32, 33).

### Statistical analysis

We performed two-tailed Mann-Whitney test to compare frequency of CD32^+^ CD4 T cells among HIV-1 uninfected, HIV-1 infected ART treated and viremic individuals. Wilcoxon signed-rank test was used to compare different subsets. Correlative analysis were done using Spearman test. These analyses were done using Prism 7.0.

Statistical analyses comparing cell surface marker expressions (non-transformed) (Fig 3A and B) in CD32^+^ CD4 T cells and subsets defined by CD32 and PD-1(i.e. CD32 vs. PD-1 quadrants) were assessed using linear mixed-effect models on the frequency of positive cells for each marker accounting for differences between patient groups (HIV-1 uninfected, HIV-1 infected ART treated and viremic individuals) with patient-level random intercepts. *P*-values were adjusted using Bonferroni correction and data analysis was performed using R statistical software and the lmer package.

## Data Availability

All relevant data are within the paper and its Supplemental Material files.

## ACKNOWLEGEMENTS

This work was supported by a grant of the Swiss National Fund. The funders had no role in study design, data collection and interpretation, or the decision to submit the work for publication.

The authors declare no competing financial interests.

We are grateful to Alex Farina, Navina Rajah, Line Leuenberger and Manon Geiser for technical assistance.

## SUPPLEMENTAL MATERIAL

**S1 Fig.**
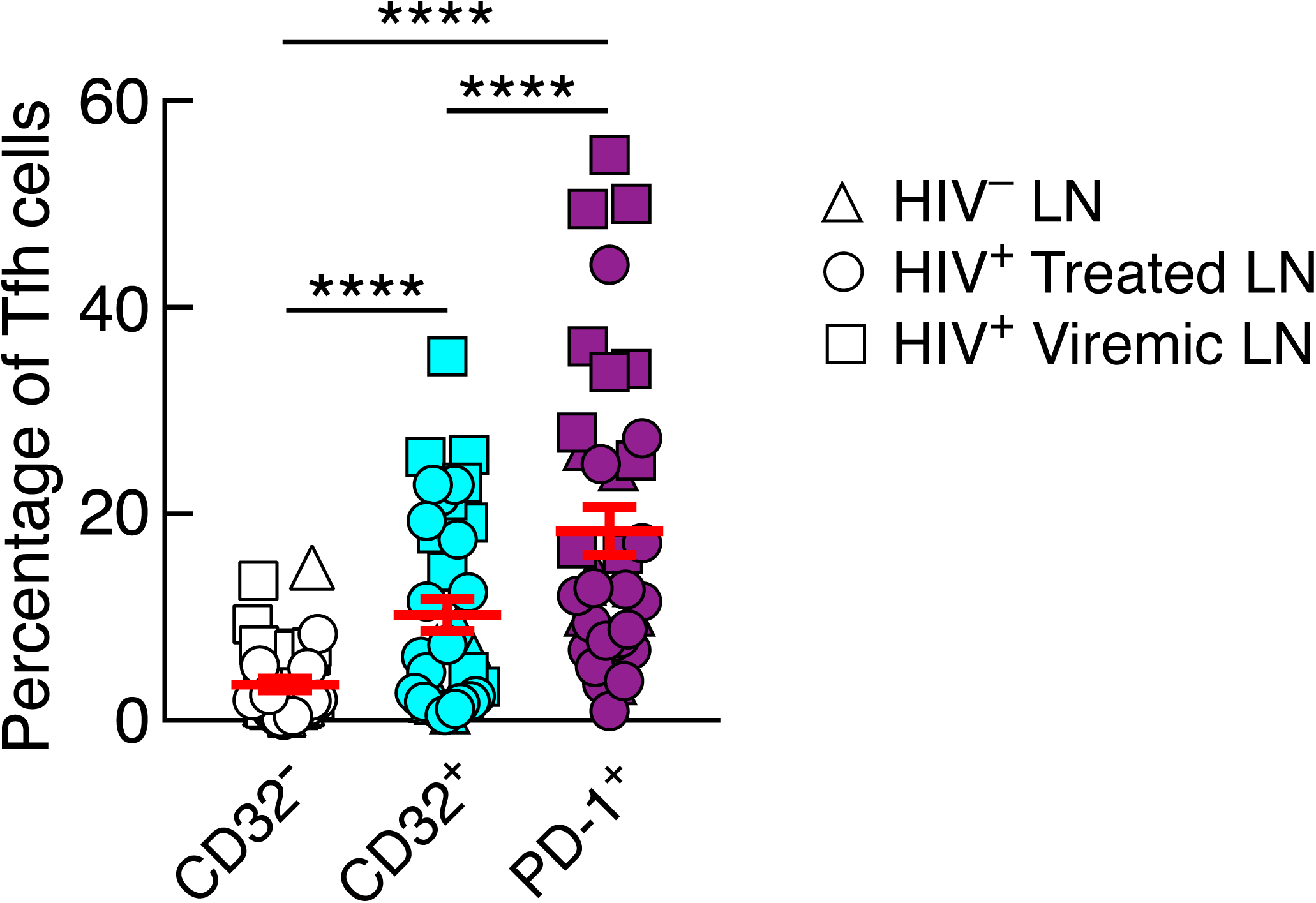
Distribution of Tfh cells within CD32-, CD32+ and PD-1+ CD4 T cell populations from lymph nodes. Triangles represent HIV-1 uninfected (N=9), round circles represent HIV-1 infected ART treated (N=19) and squares represent viremic individuals (N=10). ****P<0.0001 values were obtained by Wilcoxon signed-rank test and error bars denote mean ± S.E.M.

**S2 Fig.**
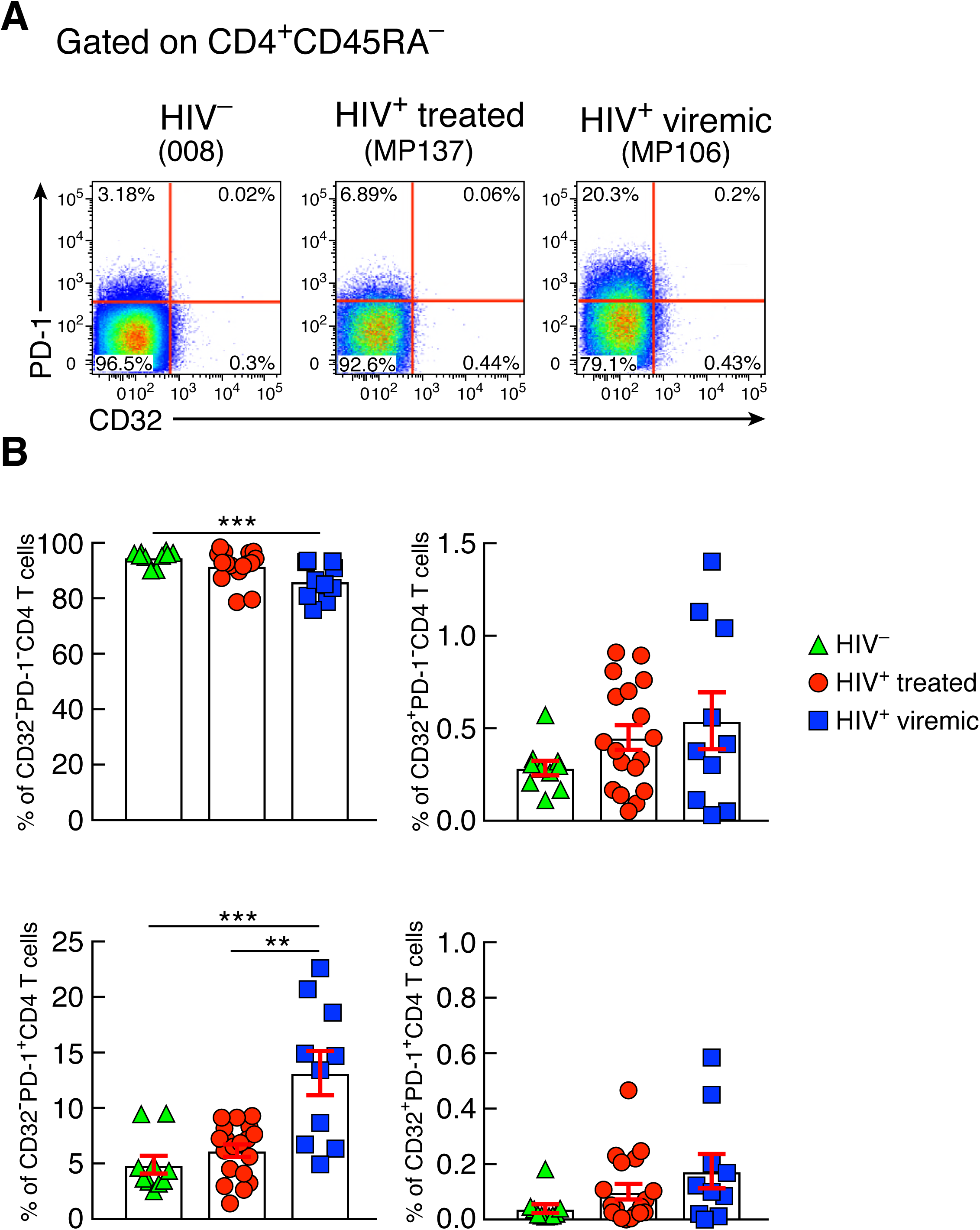
Frequencies of blood memory CD4 T cell populations on the basis of CD32 and PD-1. (A) Representative flow cytometry profiles of blood memory (CD45RA-) CD4 T cell populations expressing CD32 and/or PD-1 from representative HIV-1 uninfected, ART treated and viremic individuals.(B) Cumulative data of the frequencies of CD32-PD-1-, CD32+PD-1-, CD32-PD-1+, and CD32+PD-1+ CD4 T cell populations in blood mononuclear cells from HIV-1 uninfected (N=9), HIV-1 infected ART treated (N=16) and viremic individuals (N=10). **P<0.01 and *** P<0.001 values were obtained by Mann-Whitney test and error bars denote mean ± S.E.M

**S3 Fig.**
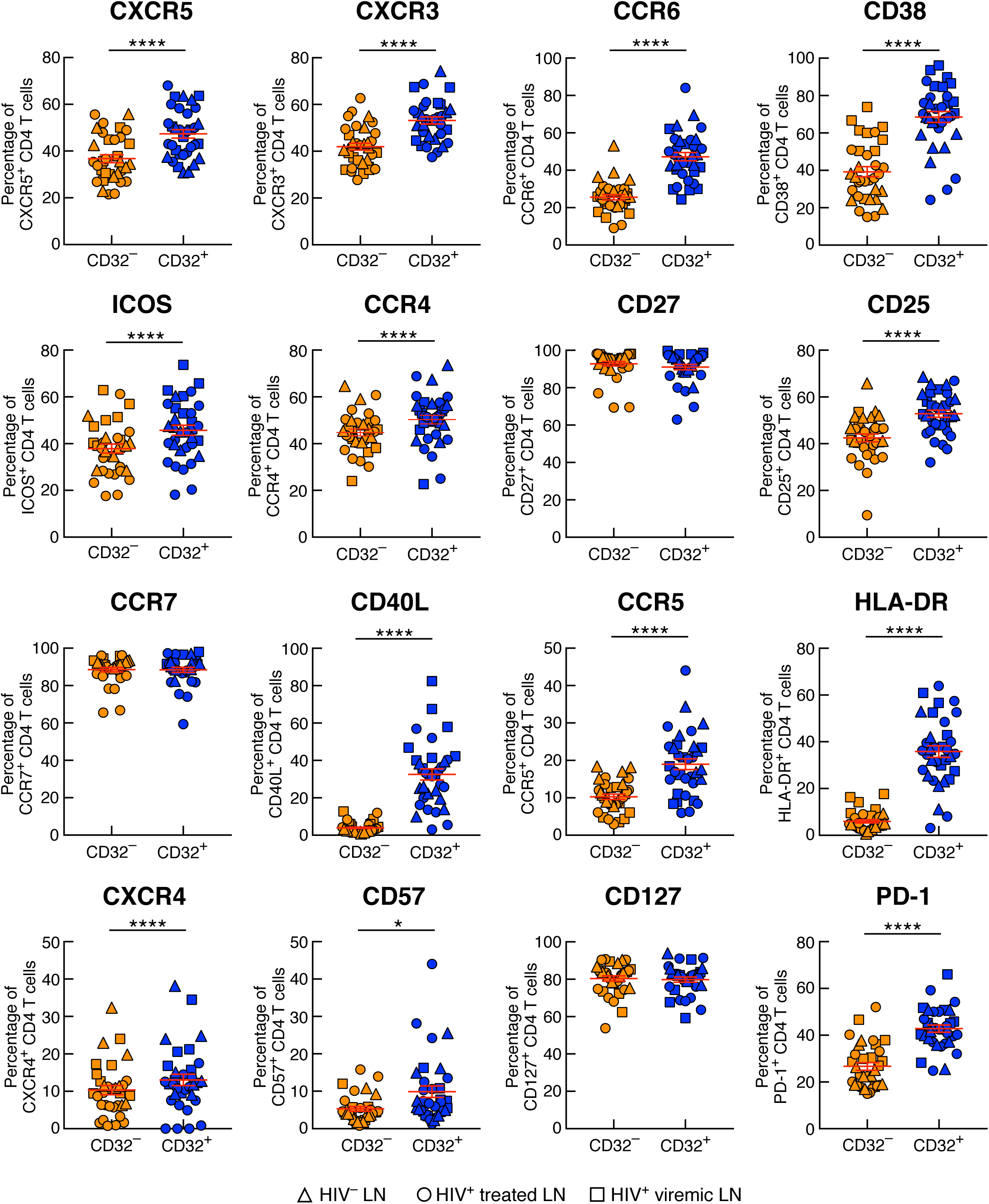
Mass cytometry profiles of CD32 expressing memory CD4 T cells from lymph nodes. Triangles represent HIV-1 uninfected (N=9), round circles represent HIV-1 infected ART treated (N=16) and squares viremic individuals (N=9). *P<0.05, ****P<0.0001 values were obtained using linear-effect mixed models with bonferroni correction for multiple testing and error bars denote mean ± S.E.M.

**S4 Fig.**
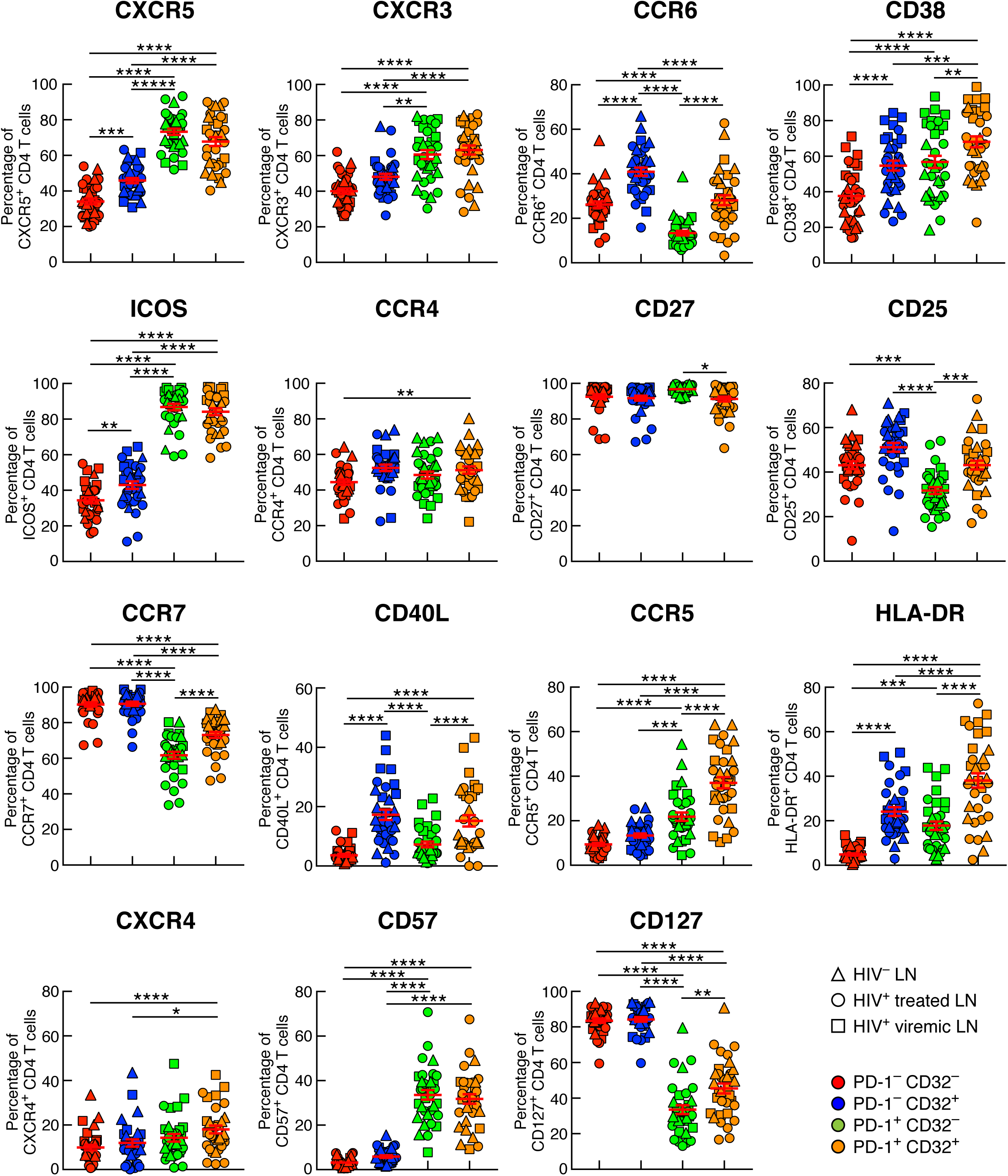
Mass cytometry analysis of LN memory CD4 T cells defined by CD32 and PD-1 expression. Triangles represent HIV-1 uninfected (N=9), round circles represent HIV-1 infected ART treated (N=16) and squares viremics individuals (N=9). *P<0.05, **P<0.01, *** P<0.001, ****P<0.0001 values were obtained using linear-effect mixed models with bonferroni correction for multiple testing and error bars denote mean ± S.E.M.

**S5 Fig.**
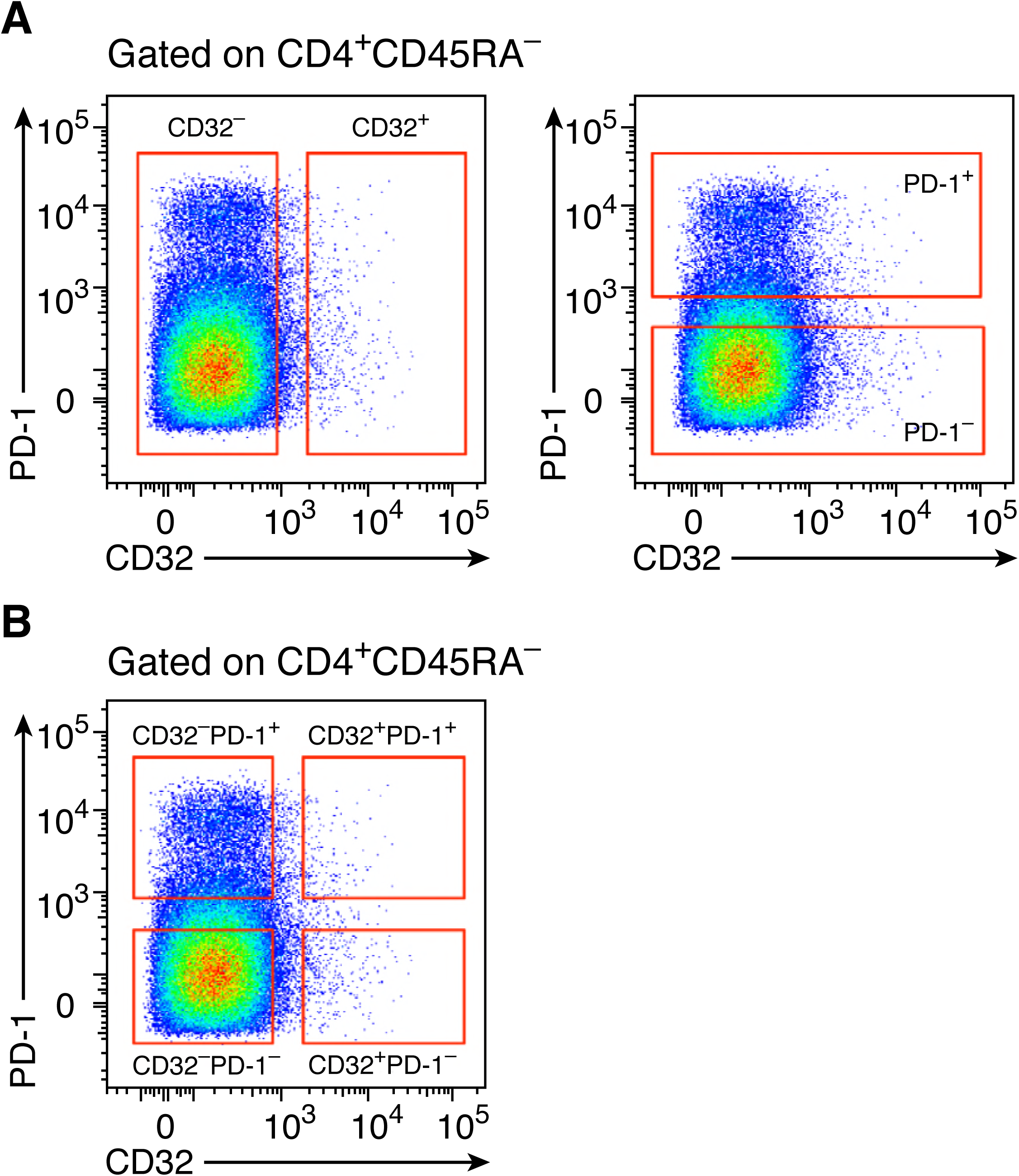
Gating strategies for cell sorting experiments. (A) gating strategy for CD32+ and CD32-(left panel) and PD-1+ and PD-1-(right panel) memory (CD45RA-) CD4 T cell populations.(B) gating strategy for CD32-PD-1-, CD32+PD-1-, CD32-PD-1+, CD32+PD-1+ memory (CD45RA-) CD4 T cell populations.

**S6 Fig.**
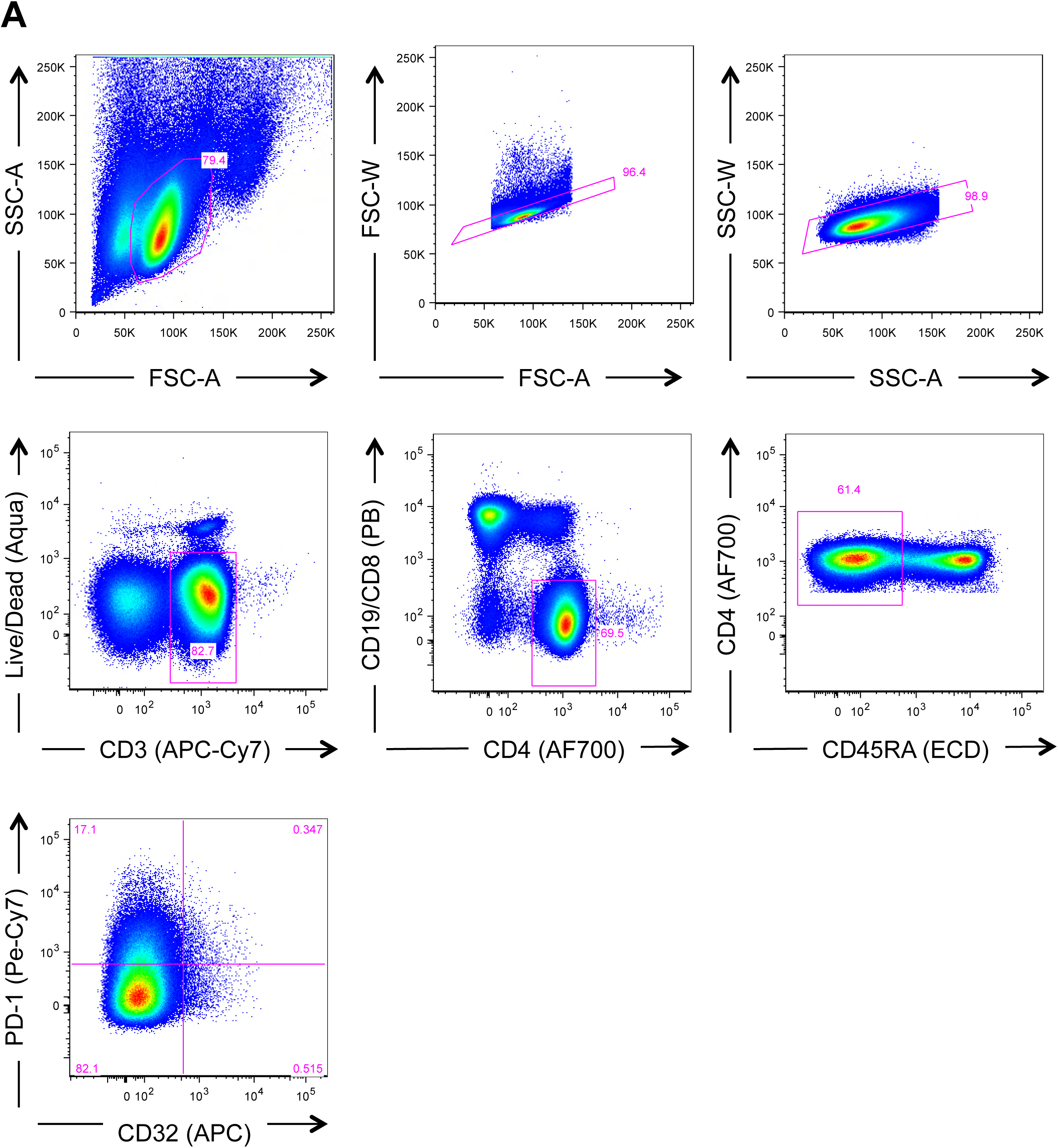
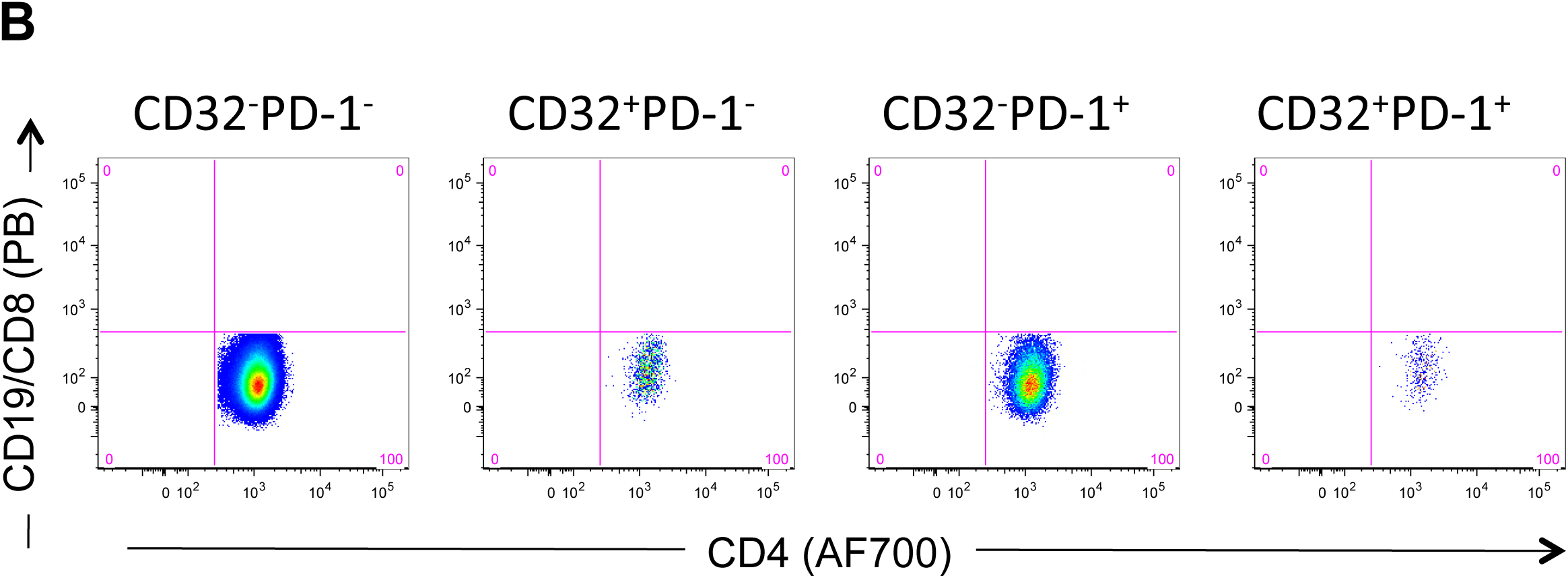
Gating strategy for flow cytometry analysis. (A) In the gating of singlet memory CD4+ T cells, lymphocytes were first gated (SSC-A vs. FSC-A) followed by singlets (FSC-A vs. FSC-W and SSC-A vs SSC-W). Lymphocytes were then gated for their uptake of the Live/Dead Aqua stain to determine live versus dead cells and for expression of CD3. On the viable CD3+ cells, we then excluded CD19 and CD8 expressing cells and gated on CD4+ cells. Low expression of CD45RA was used to gate on memory CD4+ T cells followed by the measurement of CD32 and PD-1 expression.(B) Gated CD32-PD-1-, CD32+PD-1-, CD32-PD-1+, CD32+PD-1+ cells were evaluated for the expression of CD19 and CD8 receptors.

**S1 Table.** Mass cytometry panel

| Target | Metal | Company | Clone |
| --- | --- | --- | --- |
| CD4 | 115In | Biolegend | RPA-T4 |
| CCR6 | 141Pr | Fluidigm/DVS | G034E3 |
| CD19 | 142Nd | Fluidigm/DVS | HIB19 |
| ICOS | 143Nd | Biolegend | C398.4A |
| CD8 | 145Nd | Biolegend | RPA-T8 |
| IgD | 146Nd | Fluidigm/DVS | IA6-2 |
| CD7 | 147Sm | Fluidigm/DVS | CD7-6B7 |
| CD57 | 148Nd | BD | G10F5 |
| CCR4 | 149Sm | Fluidigm/DVS | 205410 |
| CXCR5 | 153Eu | Fluidigm/DVS | RF8B2 |
| CD21 | 152Sm | Fluidigm/DVS | BL13 |
| CXCR3 | 154Sm | Biolegend | G025H7 |
| CD27 | 155Gd | Fluidigm/DVS | L128 |
| CD11c | 156Gd | Biolegend | 3.9 |
| CCR7 | 159Tb | Fluidigm/DVS | G043H7 |
| CD25 | 158Gd | Biolegend | M-A251 |
| CD14 | 160Gd | Fluidigm/DVS | M5E2 |
| CD1C | 161Dy | Biolegend | L161 |
| CD32-APC | 162Dy | Fluidigm/DVS | FUN2 |
| CD20 | 166Er | Fluidigm/DVS | 2H7 |
| CD38 | 167Er | Fluidigm/DVS | HIT2 |
| CD45RA | 169Tm | Fluidigm/DVS | HI100 |
| CD40L | 168Er | Fluidigm/DVS | CD40L |
| CD3 | 170Er | Fluidigm/DVS | UCHT1 |
| CCR5 | 171Yb | Fluidigm/DVS | NP-6G4 |
| HLA-DR | 173Yb | Fluidigm/DVS | L243 |
| PD-1 | 174Yb | Fluidigm/DVS | EH12.2H7 |
| CXCR4 | 175Lu | Fluidigm/DVS | 12G5 |
| CD127 | 176Yb | Fluidigm/DVS | A019D5 |
| CD16 | 209Bi | Fluidigm/DVS | 3G8 |
| Live/Dead | 195Pt | Fluidigm/DVS | Cell-ID |

